# Distinct signatures of calcium activity in brain pericytes

**DOI:** 10.1101/2020.07.16.207076

**Authors:** Chaim Glück, Kim David Ferrari, Annika Keller, Aiman S. Saab, Jillian L. Stobart, Bruno Weber

## Abstract

Even though pericytes have been implicated in various neurological disorders, little is known about their function and signaling pathways in the healthy brain. Here, we characterized cortical pericyte calcium dynamics using two-photon imaging of *Pdgfrβ*-CreERT2;GCaMP6s mice under anesthesia *in vivo* and in brain slices *ex vivo*. We found distinct differences between pericyte subtypes *in vivo*: Ensheathing pericytes exhibited smooth muscle cell-like calcium dynamics, while calcium signals in capillary pericytes were irregular, higher in frequency and occurred in cellular microdomains. In contrast to ensheathing pericytes, capillary pericytes retained their spontaneous calcium signals during prolonged anesthesia and in the absence of blood flow *ex vivo*. Chemogenetic activation of neurons *in vivo* and acute increase of extracellular potassium in brain slices strongly decreased calcium activity in capillary pericytes. We propose that neuronal activity-induced elevations in extracellular potassium suppress calcium activity in capillary pericytes, likely mediated by Kir2.2 and K_ATP_ channel activation.

## Introduction

The entire abluminal surface of the cerebral vasculature is covered by mural cells, namely vascular smooth muscle cells (SMC) and pericytes, which exhibit a continuum of phenotypes. Despite several attempts to categorize these heterogeneous cells in the adult mouse by morphology and molecular footprints (Hill et al., 2015, Hartmann et al., 2015a, Hartmann et al., 2015b, Vanlandewijck et al., 2018), their identities remain unclear. In general, SMCs are ring-shaped cells expressing α-smooth muscle actin (αSMA), which surround arteries and penetrating arterioles. Pericytes are embedded within the vascular basement membrane and are classically described to have a protruding “bump on a log” cell body with processes that run longitudinally along capillaries (Attwell et al., 2016). Many morphologically different subtypes of pericytes have been described (Grant et al., 2017, Uemura et al., 2020).

Pericytes regulate vascular morphogenesis and maturation as well as maintenance of the blood brain barrier (BBB, Armulik et al., 2011, Armulik et al., 2010, Daneman et al., 2010). In the context of vascular development, several signaling pathways between endothelial cells and pericytes are important; for instance platelet-derived growth factor B (PDGFB) and platelet-derived growth factor receptor beta (PDGFRβ) signaling in the recruitment of mural cells (Gaengel et al., 2009) and Angiopoietin/Tie2 signaling in angiogenesis (Teichert et al., 2017). Furthermore, CNS pericytes in the mature cortex may provide a basal tone to the vasculature and contribute to vessel stability (Berthiaume et al., 2018). In recent years, pathologies such as diabetic retinopathy, Alzheimer’s disease, circulatory failure and primary familial brain calcification have been attributed to the loss or dysfunction of pericytes (Kisler et al., 2017, Liu et al., 2017, Montagne et al., 2018, Winkler et al., 2014, Zarb et al., 2019, Nikolakopoulou et al., 2019). Yet, little is known about the function and signaling properties of pericytes in the healthy brain.

Being part of the neurovascular unit (NVU), SMCs and pericytes, positioned at the abluminal side of endothelial cells, are in close contact with astrocytes, neurons, oligodendrocytes and microglia. The NVU is involved in intricate regulatory mechanisms to tightly control blood flow (Iadecola, 2017). One of these mechanisms is functional hyperemia, which couples neural activity with changes in cerebral blood flow (CBF). Also, intrinsic vascular tone oscillations, known as vasomotion, have been observed in the brain at rest, maintaining blood “flowmotion” (Intaglietta, 1990). While it is evident that there is a relation between vasomotion and cytosolic calcium levels in SMCs (Filosa et al., 2006, Longden et al., 2016), participation of pericytes in vasomotion is currently debated (Hill et al., 2015, Hall et al., 2014, Fernandez-Klett et al., 2010, Peppiatt et al., 2006). Recently, a study in the murine olfactory bulb reported basal calcium activity in capillary pericytes and a decrease of cytosolic calcium in all mural cells along the vascular arbor as an effect of neuronal stimulation with odorants (Rungta et al., 2019). A similar observation showing a cytosolic calcium drop in pericytes was demonstrated during spreading depolarizations (Khennouf et al., 2018). However, the underlying mechanism of this calcium drop has not been identified.

In this study, we combined *in vivo* and *ex vivo* approaches to investigate calcium signaling of different pericyte subtypes in the somatosensory cortex vasculature of healthy adult mice. Since there is no consensus on the role of pericytes in blood flow control, we wondered whether morphologically different pericyte subtypes (Hartmann et al., 2015a, Grant et al., 2017) would exhibit different calcium dynamics. Furthermore, we investigated how different stimuli such as vasomodulators and neuronal activity impact the calcium dynamics of capillary pericytes.

## Results

### Two-photon imaging of *Pdgfrβ*-driven GCaMP6s in mural cells

To study calcium dynamics in mural cells, we crossed *Pdgfrβ*-CreERT2 mice (Gerl et al., 2015)) with stop-flox GCaMP6s reporter mice (Figure 1A). For localization and classification of mural cells, we defined the vessel types based on their branch order and diameter. The continuum of mural cells along the arteriovenous axis was categorized, as described earlier (Grant et al., 2017), into smooth muscle cells (SMC, 0^th^ branch), ensheathing pericytes (EP, 1^st^ to 4^th^ branch), capillary pericytes (CP, >4^th^ branch) and venular pericytes (VP) at postcapillary venules (Figure 1B and Figure 1 – figure supplement 1A-F). Noteworthy is, that in rare cases CPs interconnected vessel segments by extending processes through the parenchyma (Figure 1 – figure supplement 1D). These interconnecting pericytes were excluded from our analysis, since we do not know whether they are remnants of vessel regression (Brown, 2010) or they are a degenerative phenotype. Besides mural cells, some GCaMP6s-expressing astrocytes were sparsely detected (Figure 1B) and verified by labeling with astrocyte dye sulforhodamine 101 (SR101, Figure 1 – figure supplement 2). To avoid overlap of pericytic calcium signals by astrocytic signals, we omitted regions where a differentiation between GCaMP6s signals from astrocytes and pericytes was not possible.

**Figure 1:**
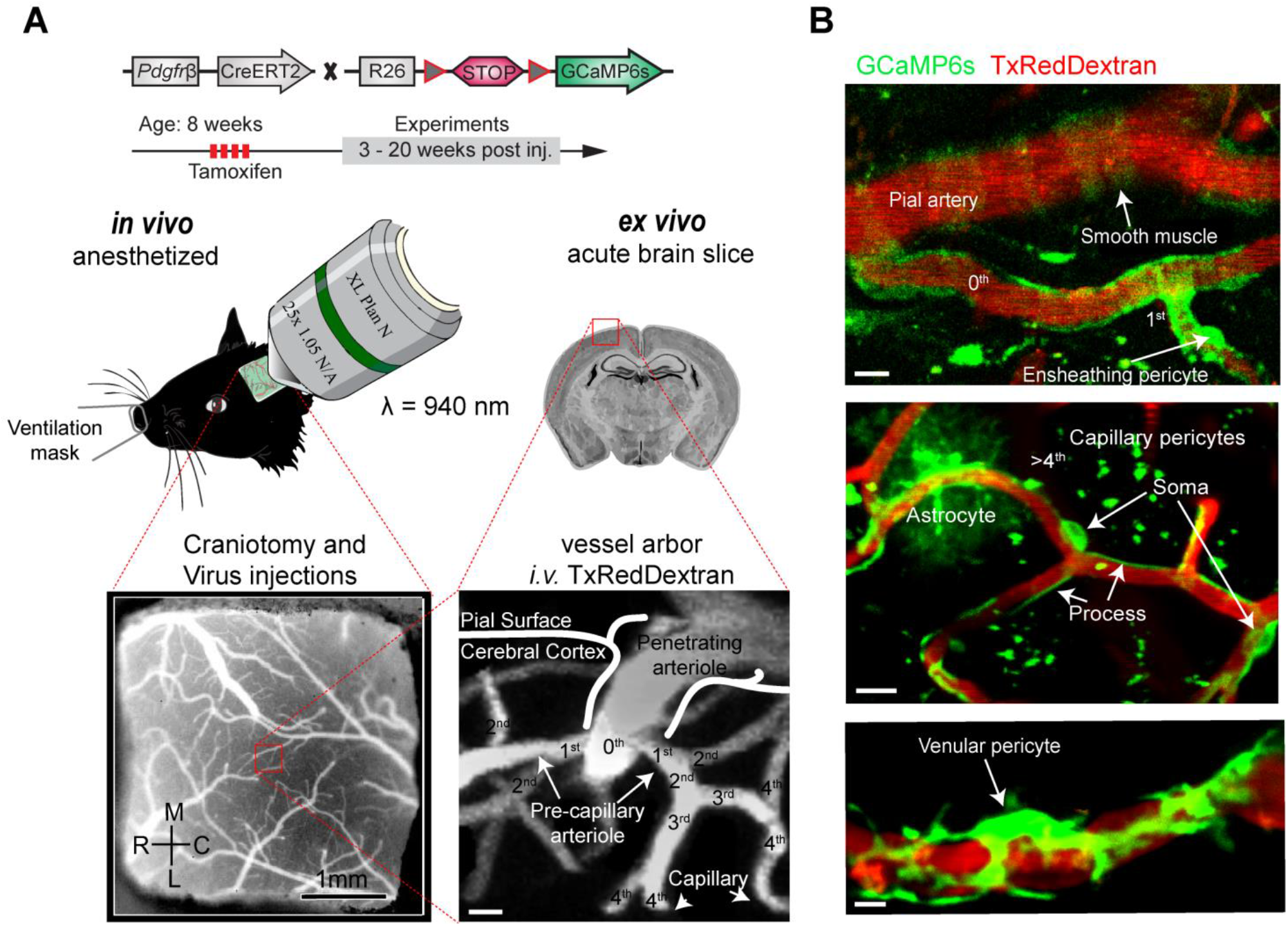
Two-photon calcium imaging of mural cells. (A) GCaMP6s expression in *Pdgfrβ*-positive cells of *Pdgfrβ*-CreERT2:R26-GCaMP6s^f/stop/f^ transgenic mice, induced by four consecutive Tamoxifen-injections. If required, adeno-associated viruses (AAV) were injected before the chronic cranial window implantation over the somatosensory cortex. Images were acquired at a wavelength of 940 nm and the vasculature was labeled via an intravenous injection (iv.) of 2.5% Texas Red Dextran (70 kDa). *In vivo* experiments were conducted with anesthetized (1.2% isoflurane) mice. For pharmacologic interventions, acute brain slices of the same mice were prepared. Z-stacks of the vessel arbor were acquired to determine the precise location of the imaged cells along the vasculature. (B) *In vivo* images of the cortical vasculature showing GCaMP6s (green) labeled mural cells. Smooth muscle cells are located on pial arteries and ensheathing pericytes on 1^st^ to 4^th^ branch order vessels. Capillary pericytes (consisting of mesh and thin stranded pericytes) are found on vessels of >4^th^ branch order. Occasionally, some astrocytes showed GCaMP6s expression. Venular pericytes reside at postcapillary venules. Scale bars: 10 μm. See Figure 1 - figure supplement 1 for mural cell identification and figure supplement 2 for GCaMP6s expressing astrocytes.

### Distinct basal calcium transients of mural cells *in vivo*

In all the mural cells described above we observed basal calcium fluctuations in somata (S) and processes (P) *in vivo* (Figure 2A, Videos 1-4). Basal calcium signals in SMCs and EPs were usually only visible during the first 30 minutes of anesthesia. Prolonged isoflurane anesthesia caused a reduction in spontaneous vasomotion, which was also reported in other studies (van Veluw et al., 2020). However, CPs continued to show basal calcium transients during the entire imaging period (under 2 hours). Visual inspection of the calcium traces suggests EPs are similar to SMCs, as both show synchronous calcium signals within somata and processes. Contrarily, CPs and VPs exhibit asynchronous calcium signals confined to microdomains along the processes (Figure 2A).

**Figure 2:**
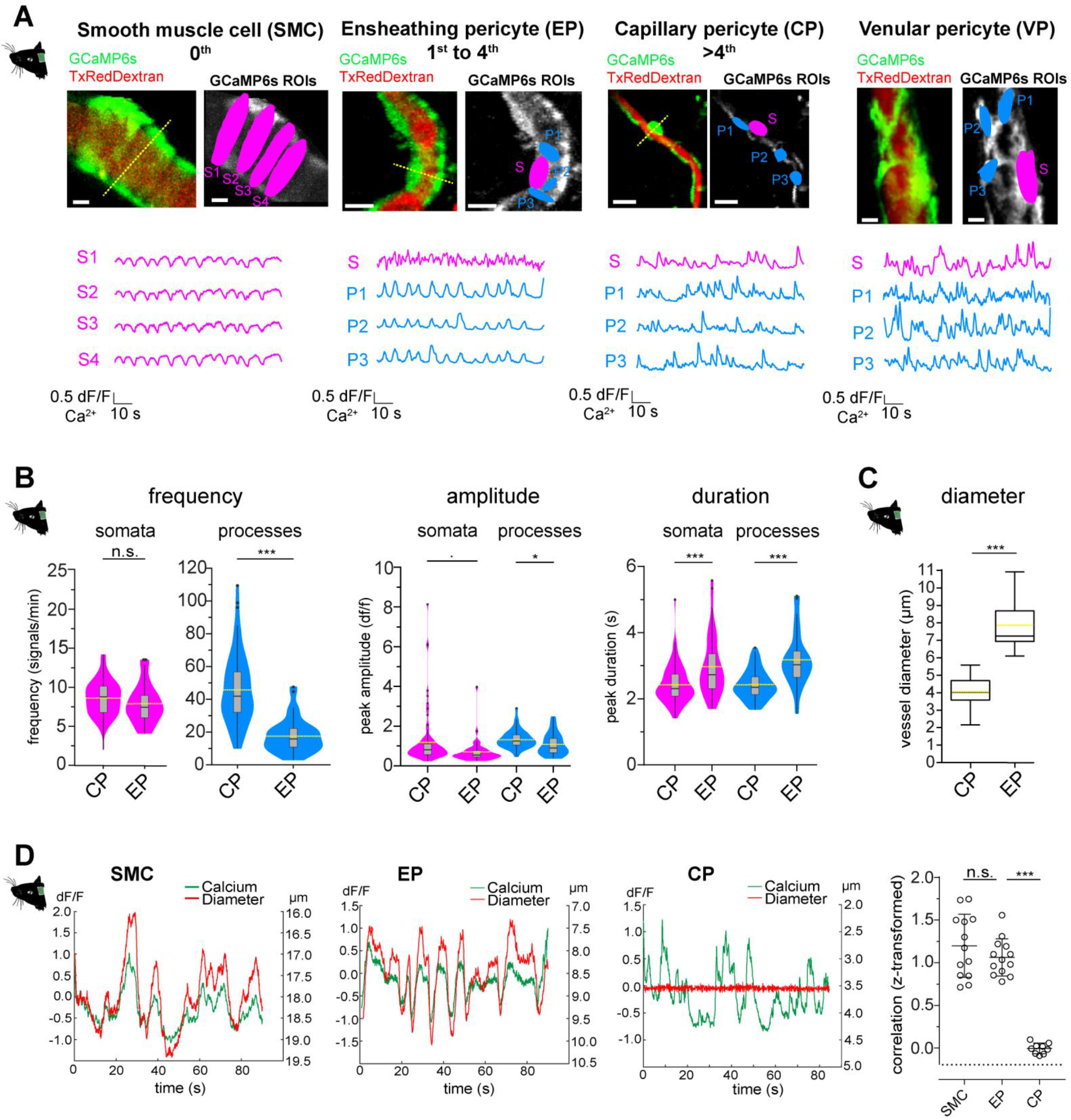
Different calcium dynamics of ensheathing and capillary pericytes. (A) Representative images of SMCs, EP, CP and VP *in vivo*. Regions of interest (ROIs) for somata (S, magenta) were hand selected. ROIs for processes (P1-3, blue) were found in an unbiased way, employing a MATLAB-based algorithm (see Figure 2 – figure supplement 1). Below the images are the normalized time traces of calcium signals of GCaMP6s fluorescence extracted from the corresponding example ROIs. Vessel diameters were measured with linescans across somata (indicated by the dashed yellow lines). Scale bars: 10 μm. (B) Comparison of quantified baseline calcium transient parameters: frequency (signals per minute), peak amplitude (dF/F) and duration (s) between CP and EP somata and processes *in vivo*, shown as violin/box plots (Tukey whiskers). The dashed yellow lines indicate the mean values of the respective parameters. Statistics were calculated using linear mixed-effects models and Tukey post hoc tests, CP: N=25, n=93, EP: N=16, n=44. Frequency, S: *P* = 0.162, P: *P* < 0.001; amplitude, S: *P* = 0.051, P: *P* = 0.011; duration, S: *P* < 0.001, P: *P* < 0.001. (C) Comparison of the inner vessel diameter, measured at somata of capillary (n=54) and ensheathing (n=23) pericytes, shown in a box plot (Tukey whiskers). Unpaired two-tailed t-test, t(75) = 15.21, *P* < 0.001, the dashed yellow lines indicate the mean values. (D) Plots depicting the relation between cytosolic calcium (green trace) and vessel diameter (red trace, note that the left y-axis for diameter is inverted) measured simultaneously with line scans along the soma, for SMC, EP and CP. Traces were centered on the average diameter of the vessel. On the far-right comparison of the correlations (Fisher z-transformed) between calcium signals and changes in vessel diameter (inverted). Unpaired two-tailed t-tests were performed to compare groups. Individual values with mean (SD) are shown. SMC (n = 13), EP (n = 13), CP (n = 9). n.s. indicates not significant, t(24) = 1.139, *P* = 0.27; t(20) = 14.21, *P* < 0.001. See Figure 2 – figure supplement 1 for automated ROI detection and figure supplement 2 for power spectral analysis of EPs. Source data: Figure 2 – Source Data 1.

To compare the calcium dynamics between EPs and CPs, we measured frequency, amplitude and duration of calcium transients using semi-automated image analysis as described and employed earlier by our lab (Zuend et al., 2020, Stobart et al., 2018). Regions of interest (ROI) for pericyte somata (S) were selected by hand and ROIs for pericyte processes (P) were automatically detected with an unbiased algorithm (Ellefsen et al., 2014) implemented in our MATLAB toolbox, CHIPS (Barrett et al., 2018). EPs displayed a smaller number of calcium signals/min in somata and processes that were also of longer duration (frequency, S: 7.8 (2.5) signals/min and P: 18.9 (9.4) signals/min; duration, S: 3.0 (0.9) s and P: 3.2 (0.8) s), compared to CPs (frequency, S: 8.5 (2.2) signals/min and P: 45.7 (19.3) signals/min; duration, S: 2.4 (0.6) s and P: 2.4 (0.4) s. Values are mean (SD). Figure 2B).

In line with previous reports (Hartmann et al., 2015a, Grant et al., 2017), we found EPs on vessels of 1^st^ to 4^th^ branch order, with an average inner diameter of 7.9 (1.5) μm, n=23, whereas CPs were found on capillaries (>4^th^ branch order) with an average diameter of 4.1 (0.7) μm, n=54 (Values are mean (SD). Figure 2C).

Calcium signals in SMCs during spontaneous vasomotion are inversely correlated to vessel diameter (transformed r _Fisher z_ = 1.20 (0.37), n=13), as evaluated by cross-correlation analysis (Figure 2D), and reported in a previous study (Hill et al., 2015). We observed a similar behaviour in EPs (transformed r _Fisher z_ = 1.06 (0.22), n=13; Figure 2D). Moreover, power spectral analysis of the EPs calcium signals shows a distinct peak at 0.1 Hz (Figure 2 – figure supplement 2). This is in line with spontaneous vasomotion described in SMCs (van Veluw et al., 2020, Mateo et al., 2017). In accordance with earlier studies (Pabelick et al., 2001), we found that the vessel diameter change follows the calcium change in EPs by an average delay of 300 ms, suggesting a calcium-dependent contraction mechanism. In contrast, there is no relationship between calcium signals in CPs and vessel diameter (transformed r _Fisher z_ = −0.005 (0.06), n = 9; Figure 2D).

Thus, EPs and CPs display different basal calcium dynamics, which besides their location in the vascular network, morphology and αSMA-expression, can be used as a further measure to differentiate between these pericyte subtypes.

### Persisting calcium signals in CPs *ex vivo*

To further investigate the differences in calcium signals between EPs and CPs, we prepared acute brain slices for *ex vivo* pharmacological probing. Blood plasma was stained with an iv. injection (100 μL, 2.5%) of 70 kDa Texas Red Dextran via the tail vein prior to the slice preparation. The fluorophore remains in the vasculature for several hours for easy capillary identification and further serves as a control for motion artefacts. Orientation in the slice was obtained by following a penetrating artery from the slice surface to its branches.

Calcium signals of EPs were greatly reduced *ex vivo*. Comparing the signal frequency between *in vivo* and *ex vivo,* there was a more than twofold reduction in calcium activity in both somata and processes (*in vivo*, S: 7.8 (2.5) and P: 18.9 (9.4); *ex vivo*, S: 3.1 (1.6) and P: 7.3 (5.6). Values are mean (SD) signals/min. Figure 3A). In many cases (Figure 3B), arteries and arterioles were either collapsed due to loss of tone or intraluminal pressure in the slice preparation or were constricted by possibly dead SMCs or EPs (no detectable calcium signal).

**Figure 3:**
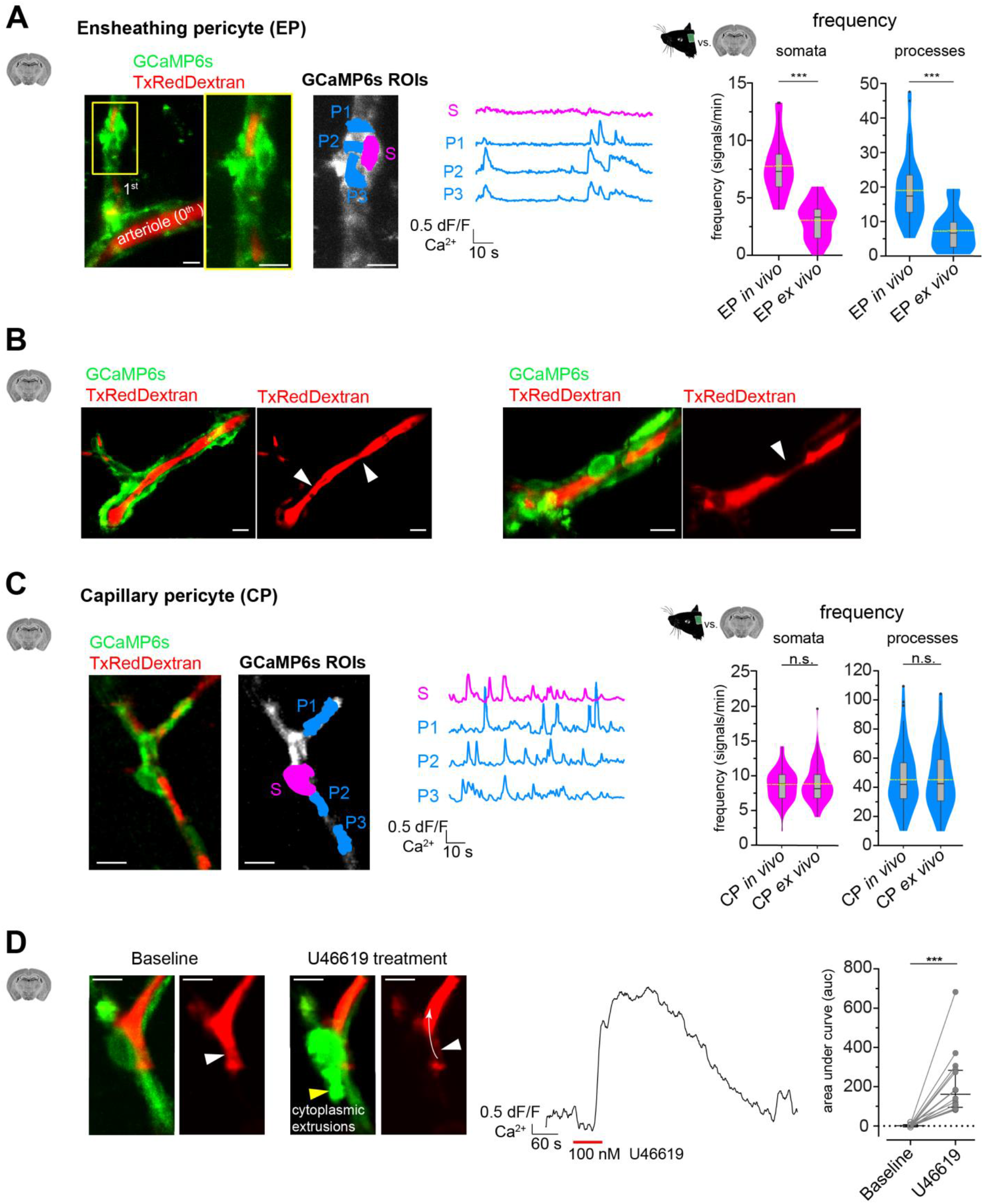
Calcium dynamics in the absence of blood flow *ex vivo* is more affected in EPs than CPs. (A) Ensheathing pericytes *ex vivo* can be localized similarly to those *in vivo* by following the vessel branches of a pial artery lying on the surface of the brain slice. On the left, two-photon microscopy image of a precapillary arteriole and its adjacent 1^st^ order branch, where an ensheathing pericyte is located. The yellow box shows a magnified image of the ensheathing pericyte. In the GCaMP6s channel, ROIs for soma (S, in magenta) and processes (P1-3, in blue) are shown. In the center are the respective normalized traces of calcium signals. On the right are violin/box plots depicting the quantified calcium signal frequency of ensheathing pericytes *ex vivo* compared to the previously (Figure 2B) determined calcium signal frequency *in vivo* for somata and processes. The dashed yellow line indicates the mean value. Statistics were calculated using linear mixed-effects models and Tukey post hoc tests. *In vivo*: N=16, n=44, *ex vivo*: N=10, n=22. S: *P* < 0.001; P: *P* < 0.001. (B) On the left, vessels in *ex vivo* acute brain slices tend to collapse (see white arrowheads). Vessel segments with ensheathing pericytes (on the right) are often to be found constricted at the soma (white arrowhead). (C) On the left, representative image of a capillary pericyte *ex vivo*. In the GCaMP6s channel, ROIs for soma (S, in magenta) and processes (P1-3, in blue) are shown. In the center are the respective normalized traces of calcium signals. On the right are violin/box plots depicting the quantified calcium signal frequency of capillary pericytes *ex vivo* compared to the previously (Figure 2B) determined calcium signal frequency *in vivo* for somata and processes. The dashed yellow lines indicate the mean values. Statistics were calculated using linear mixed-effects models and Tukey post hoc tests. *In vivo*: N=25, n=93, *ex vivo*: N=52, n=106. S: *P* = 0.552; P: *P* = 0.806. (D) On the left, application of a thromboxane A_2_ receptor agonist U46619 (100 nM) leads to a massive calcium response in capillary pericytes and is accompanied by cytoplasmic extrusions (yellow arrowhead). The corresponding vessel image shows that blood plasma is moved away (white arrow indicates the direction of movement) from the site where the soma is located in the center the normalized trace of the calcium response of the whole cell is shown. On the right, quantification of the area under the curve (auc) comparing baseline to U46619 treatment is shown. Data represents individual cells, median and interquartile ranges. Statistics were calculated using a two-tailed Wilcoxon matched-pairs signed rank test. N=5, n=16, *P* < 0.001. Scale bars: 10 μm. n.s. indicates not significant, \**P* < 0.05, \*\**P* < 0.01, \*\*\**P* < 0.001. See Figure 3 – figure supplement 1 for further calcium signal comparison between *in vivo* and *ex vivo* CPs. See Figure 3 – figure supplement 2 for calcium signal comparison between *in vivo* and *ex vivo* VPs. Source data: Figure 3 – Source Data 1.

Surprisingly, CPs in *ex vivo* brain slices retained their highly frequent calcium transients, which were also confined to microdomains (Figure 3C). The comparison of basal calcium transients between *in vivo* and *ex vivo* CPs did not reveal a difference with respect to their signal frequency (*in vivo*, S: 8.5 (2.2) and P: 45.7 (19.3); *ex vivo*, S: 8.7 (2.6) and P: 44.8 (19.8). Values are mean (SD) signals/min). However, the signal amplitude and the duration differed between *in vivo* and *ex vivo* (amplitude: *in vivo*, S: 1.1 (1.3) df/f and P: 1.3 (0.4) df/f; *ex vivo*, S: 1.8 (1.8) df/f and P: 1.8 (0.7) df/f; duration: *in vivo*, S: 2.4 (0.6) s and P: 2.4 (0.4) s; *ex vivo*, S: 2.1 (0.5) s and P: 2.2 (0.4) s. Values are mean (SD). Figure 3 – figure supplement 1). Nonetheless, due to the similar calcium frequency dynamics *in vivo* and *ex vivo*, we continued to use acute brain slices, with a focus on CPs, to probe pharmacologic agents in an attempt to understand how calcium signals are regulated.

Worthy of note is, that VPs also retained their calcium dynamics *ex vivo* with no evident change compared to *in vivo* (*in vivo*, S: 8.2 (0.9) signals/min, 1.2 (1.2) df/f, 2.1 (0.3) s and P: 47.2 (26.4) signals/min, 1.3 (0.7) df/f, 2.3 (0.4) s; *ex vivo*, S: 8.4 (3.1) signals/min, 1.9 (3.0) df/f, 2.0 (0.4) s and P: 35.8 (17.6) signals/min, 1.7 (0.6) df/f, 2.4 (0.5) s. Values are mean (SD). Figure 3 – figure supplement 2).

The application of the thromboxane A_2_ receptor agonist U46619 (100 nM) to the slice preparation is often used to prevent the collapse of wider calibre vessels by generating an artificial vascular tone (Mishra et al., 2014, Brown et al., 2002). However, we observed a massive calcium response in CPs when U46619 was included in the superfusate (Figure 3D, Video 5). This overt calcium response in CPs was accompanied by cytoplasmic extrusions, suggesting that U46619 might be toxic for pericytes. In addition, blood plasma inside the vessel was pushed away from the region where the CP soma was located (Figure 3D). Moreover, the whole vasculature responded to the global reactivity of U46619. For this reason, we omitted the use of any preconstricting agent in our slice preparations.

### CP calcium events are evoked by vasomodulators

Next, we probed CPs for their responsiveness to different vasomodulators, such as G-protein-coupled receptor (GPCR) agonists Endothelin-1 (100 nM), UDP-Glucose (100 μM) and ATP (100 μM). All agonists elevated cytosolic calcium levels, suggesting a responsiveness of CPs to endothelial or astrocytic-derived factors (Figures 4A-C).

**Figure 4:**
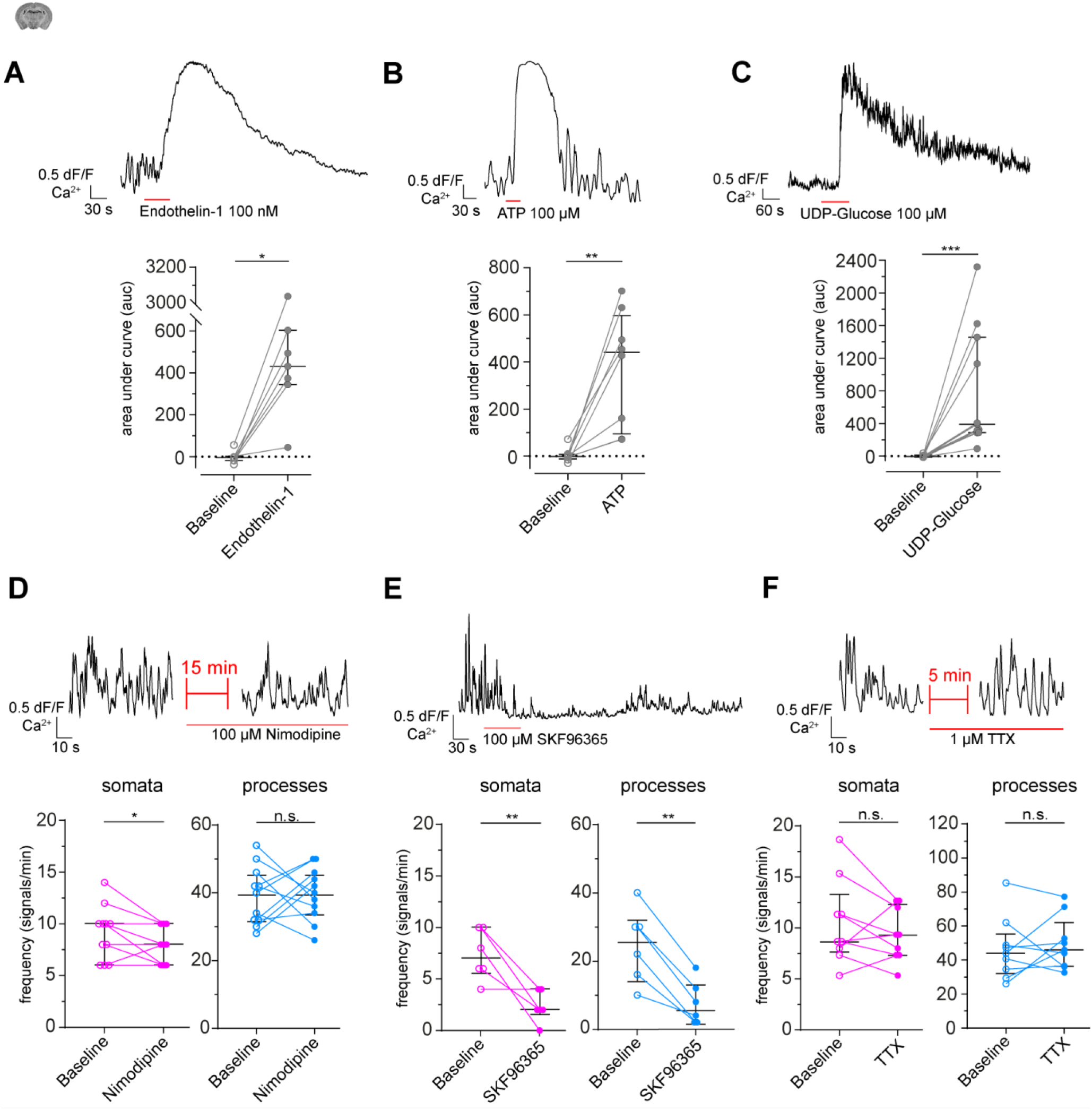
Modulation of capillary pericyte calcium signals *ex vivo*. (A-C) Calcium responses to the application of vasomodulators Endothelin-1 (100 nM), ATP (100 μM) and UDP-Glucose (100 μM). On top are representative normalized calcium signal traces and below are quantifications of the area under the curve (auc), comparing baseline to the respective treatment. Data represents individual cells, median and interquartile ranges. Statistics were calculated using Wilcoxon matched-pairs signed rank tests. Endothelin-1: N=3, n=7, *P* = 0.02; ATP: N=4, n=8, *P* = 0.008; UDP-Glucose: N=4, n=11, *P* < 0.001. (D) Calcium response to application of the L-type voltage-gated calcium channel (L-type VGCC) blocker Nimodipine (100 μM). On top is a representative normalized calcium signal trace and below is the quantification of the signal frequency in somata and processes, comparing baseline to the Nimodipine treatment. Nimodipine was infused 15 min prior to data collection. Data represents individual cells, median and interquartile ranges. Statistics were calculated using two-tailed paired t-tests, N=4, n=11. S: t(10) = 2.39, *P* = 0.04; P: t(10) = 0.1495, *P* = 0.88. (E) Calcium response to application of the TRPC channel blocker SKF96365 (100 μM). On top is a representative normalized calcium signal trace and below is the quantification of the signal frequency in somata and processes, comparing baseline to the SKF96365 treatment. Data represents individual cells, median and interquartile ranges. Statistics were calculated using two-tailed paired t-tests, N=3, n=6. S: t(5) = 4.038, *P* = 0.01; P: t(5) = 6.755, *P* = 0.001. (F) Calcium response to application of TTX (1 μM). On top is a representative normalized calcium signal trace and below is the quantification of the signal frequency in somata and processes, comparing baseline to the TTX treatment. TTX was infused 5 min prior to data collection. Data represents individual cells, median and interquartile ranges. Statistics were calculated using two-tailed paired t-tests, N=3, n=9. S: t(8) = 1.113, *P* = 0.2979; P: t(8) = 0.5425, *P* = 0.6. The red lines below the traces indicate the addition time of the respective drug. n.s. indicates not significant, \**P* < 0.05, \*\**P* < 0.01,\*\*\**P* < 0.001. Source data: Figure 4 – Source Data 1.

Since the investigated GPCRs are mainly acting via phospholipase C and subsequent IP_3_-mediated calcium release from internal stores (Wynne et al., 2009, Lazarowski and Harden, 2015, Abbracchio et al., 2006), we continued to examine the involvement of ion channels, which have been shown to affect cytosolic calcium levels in SMCs (Hill-Eubanks et al., 2011). Application of Nimodipine (100 μM), a blocker of L-type voltage-gated calcium channels (VGCC) involved in SMC contractions, led to a moderate reduction of basal calcium transients in CP somata but not processes (baseline, S: 9.1 (2.6), P: 39.1 (8.6); Nimodipine, S: 7.6 (1.7), P: 39.6 (7.8). Values are mean (SD) signals/min. Figure 4D). However, a transient receptor potential channel (TRPC) blocker (Earley and Brayden, 2015), SKF96365 (100 μM), reduced the calcium transient frequency in both somata and processes (baseline, S: 7.3 (2.4), P: 24.7 (10.9); SKF96365, S: 2.3 (1.5), P: 7.7 (6.4). Values are mean (SD) signals/min. Figure 4E), showing that CPs need extracellular calcium to maintain their calcium fluctuations.

We were then interested to find out whether neurons could have an impact on the calcium dynamics of CPs. We applied tetrodotoxin (TTX), a voltage-gated sodium channel blocker, to dampen neuronal activity (Zonta et al., 2003). Nonetheless, CP calcium signals were not affected (baseline, S: 10.5 (4.2), P: 46.4 (18.2); TTX, S: 9.3 (2.6), P: 49.6 (15.6). Values are mean (SD) signals/min. Figure 4F).

### Potassium released by neuronal stimulation leads to a calcium signal drop in CPs

Neurons have a low basal activity in acute brain slices (Ballanyi and Ruangkittisakul, 2009), so we sought to specifically stimulate cortical neurons *in vivo*. We used chemogenetics (Roth, 2016), in which neurons – in this case transduced to express hM3D(Gq)-DREADD (designer receptors exclusively activated by designer drugs) – were activated with 30 μg/kg iv. clozapine (Jendryka et al., 2019). This specific neuron activation led to a drop in CP calcium signals (baseline, S: 7.0 (2.6), P: 38.7 (18.1); Clozapine, S: 0.0 (0.0), P: 2.0 (2.8). Values are mean (SD) signals/min. Figure 5A, Video 6). Non-specific effects caused by clozapine on pericytic calcium signals are unlikely because CPs outside the area of DREADD-expressing neurons retained their basal calcium dynamics.

**Figure 5:**
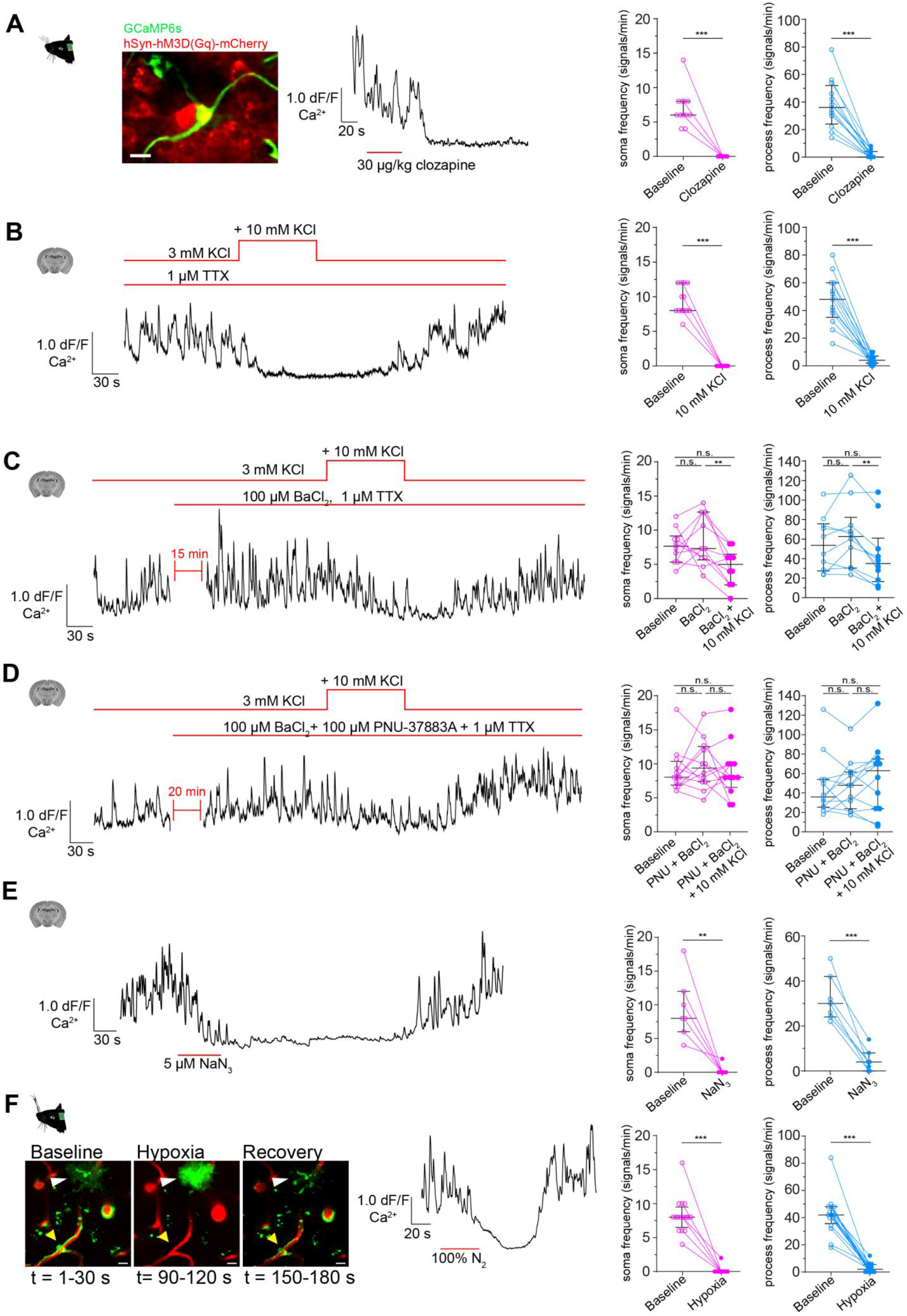
Capillary pericytes sense potassium released by activated neurons. (A) Calcium response of CPs to hM3D(Gq) chemogenetic activation of neurons *in vivo*. The image (on the left) shows a CP and neurons, expressing hM3D(Gq)-mCherry. A representative normalized calcium signal trace is shown (in the center) and the calcium signal frequency is quantified, comparing baseline to treatment (on the right). Data represents individual cells, median and interquartile ranges. Statistics were calculated using a Wilcoxon matched-pairs signed rank test (for S) and a two-tailed paired t-test (for P), N=4, n=12. S: *P* < 0.001; P: t(11) = 7.453, *P* < 0.001. Scale bar = 10 μm. (B) Calcium response of CPs to a 10 mM rise of extracellular potassium in the presence of TTX (1 μM) *ex vivo*. A representative normalized calcium signal trace (left) and the quantification of the calcium signal frequency, comparing baseline to treatment (right) are shown. Data represents individual cells, median and interquartile ranges. Statistics were calculated using a Wilcoxon matched-pairs signed rank test (for S) and a two-tailed paired t-test (for P), N=4, n=13. S: *P* < 0.001; P: t(12) = 8.516, *P* < 0.001. (C) Calcium response of CPs to a 10 mM rise of extracellular potassium in the presence of TTX (1 μM) and Kir 2 channel blocker BaCl_2_ (100 μM) *ex vivo*. A representative normalized calcium signal trace is shown on the left and on the right, the calcium signal frequency is quantified, comparing baseline to pre-treatment (TTX + BaCl_2_) and additional potassium treatment. Data represents individual cells, median and interquartile ranges. Statistics were calculated using a one-way ANOVA, N=4, n=10. S: F(1.796, 16.17) = 5.728, *P* = 0.02, *P*(baseline vs. BaCl_2_) = 0.82, *P*(baseline vs. BaCl_2_ + KCl) = 0.10, *P*(BaCl_2_ vs. BaCl_2_ + KCl) = 0.009; P: F(1.542, 13.88) = 7.137, *P* = 0.01, *P*(baseline vs. BaCl_2_) = 0.31, *P*(baseline vs. BaCl_2_ + KCl) = 0.22, *P*(BaCl_2_ vs. BaCl_2_ + KCl) = 0.002. (D) Calcium response of CPs to a 10 mM rise of extracellular potassium in the presence of TTX (1 μM), Kir 2 channel blocker BaCl_2_ (100 μM) and K_ATP_ channel blocker PNU-37883A (100 μM) *ex vivo*. A representative normalized calcium signal trace is shown on the left. On the right, the calcium signal frequency is quantified, comparing baseline to pre-treatment (TTX + BaCl_2_ + PNU-37883A) and additional potassium treatment. Data represents individual cells, median and interquartile ranges. Statistics were calculated using a one-way ANOVA, N=4, n=12. S: F(1.395, 15.35) = 0.2248, *P* = 0.72, *P*(baseline vs. PNU + BaCl_2_) = 0.76, *P*(baseline vs. PNU + BaCl_2_ + KCl) = 0.99, *P*(PNU + BaCl_2_ vs. PNU + BaCl_2_ + KCl) = 0.77; P: F(1.683, 18.51) = 0.9687, *P* = 0.38, *P*(baseline vs. PNU + BaCl_2_) = 0.96, *P*(baseline vs. PNU + BaCl_2_ + KCl) = 0.54, *P*(PNU + BaCl_2_ vs. PNU + BaCl_2_ + KCl) = 0.44. (E) Calcium response of CPs to NaN_3_ (5 μM) *ex vivo*. A representative normalized calcium signal trace is shown on the left and the quantification of the calcium signal frequency, comparing baseline to NaN3 treatment is shown on the right. Data represents individual cells, median and interquartile ranges. Statistics were calculated using two-tailed paired t-tests, N=3, n=7. S: t(6) = 4.824, *P* = 0.003; P: t(6) = 7.621, *P* < 0.001. (F) Calcium response of CPs to an acute hypoxia insult *in vivo*. On the left, two-photon images show the time course and GCaMP6s fluorescence of the hypoxic insult (yellow arrowhead: capillary pericyte; white arrowhead: astrocyte). Center, a representative normalized calcium signal trace is shown. The calcium signal frequency is quantified on the right. Baseline is compared to hypoxic intervention. Data represents individual cells, median and interquartile ranges. Statistics were calculated using Wilcoxon matched-pairs signed rank tests, N=4, n=16. S: *P* < 0.001; P: *P* < 0.001. The red lines indicate the addition time of the respective drug. Source data: Figure 5 – Source Data 1.

This observation coincides with a previous report in the olfactory bulb, in which pericytes reacted to odorant stimulation with a decrease in calcium transients (Rungta et al., 2018).

When stimulated, neurons release potassium into the extracellular space (Somjen, 1979, Heinemann and Lux, 1977). This potassium is sensed by SMCs, causing the suppression of calcium oscillations (Filosa et al., 2006). To test whether calcium signals also drop in CPs when extracellular potassium is increased, we temporarily raised the extracellular concentration of potassium (in the form of KCl) by 10 mM. Indeed, CPs reacted with a calcium signal drop in both somata and processes (baseline, S: 9.4 (2.1), P: 47.5 (17.8); KCl, S: 0.0 (0.0), P: 4.3 (3.4). Values are mean (SD) signals/min. Figure 5B). To exclude any secondary neurotransmitter effects, synaptic activity was blocked with TTX (1 μM). TTX alone did not influence the calcium dynamics of CPs (Figure 4F).

In SMCs, Kir2 channels are the main drivers for potassium sensing (Longden and Nelson, 2015). Single cell transcriptomic analysis revealed a high abundance of Kir2.2 (*Kcnj12*) transcripts in pericytes too (Vanlandewijck et al., 2018). Therefore, it is plausible that a similar mode of potassium sensing occurs in pericytes. We antagonized Kir2 channels with BaCl_2_ (100 μM), and interestingly, we saw a less pronounced decrease in CP calcium signals in response to KCl stimulation in the presence of BaCl_2_. (baseline, S: 7.7 (2.5), P: 55.1 (27.9); BaCl_2_, S: 8.5 (3.7), P: 63.5 (33.2); BaCl_2_ + KCl, S: 4.6 (2.7), P: 43.2 (33.2). Values are mean (SD) signals/min. Figure 5D).

Furthermore, an ATP-sensitive potassium channel (Nichols, 2006) composed of Kir6.1 (*Kcnj8*) and its associated sulfonylurea receptor 2 (SUR2, *Abcc9*) has been suggested as a marker for brain pericytes (Bondjers et al., 2006, Vanlandewijck et al., 2018). To see, whether this channel is involved in the potassium-mediated calcium signal decrease in CPs, we applied the Kir6.1 specific antagonist (Teramoto, 2006) PNU-37883A (100 μM), together with the Kir2 channel blocker BaCl_2_. This prevented a decrease in calcium signals in both CP somata and processes, when extracellular potassium was increased by 10 mM (baseline, S: 9.1 (3.2), P: 46.9 (31.1); PNU + BaCl_2_, S: 9.8 (3.6), P: 48.4 (25.5); PNU + BaCl_2_ + KCl, S: 9.0 (4.0), P: 55.0 (36.4). Values are mean (SD) signals/min. Figure 5D).

Next, we investigated the possible involvement of K_ATP_ channels in the regulation of calcium signaling in CPs. Lowering the cellular [ATP]/[ADP] ratio by blocking mitochondrial activity (causing ischemia) with sodium azide (NaN_3_, 5 μM) in *ex vivo* brain slices should result in K_ATP_ channel activation (Dart and Standen, 1995, Tinker et al., 2018, Reiner et al., 1990, Lerchundi et al., 2019). Indeed, CP calcium signals decreased in response to NaN_3_ (baseline, S: 9.4 (4.6), P: 32.3 (10.2); NaN_3_, S: 0.3 (0.8), P: 4.6 (5.0). Values are mean (SD) signals/min. Figure 5E). We mimicked a shortage in respiratory supply *in vivo* by creating an acute hypoxic insult, whereby oxygen in the anesthesia gas mixture was transiently replaced with pure nitrogen. In a similar manner to the *ex vivo* chemical ischemia response, CPs responded to hypoxia with a reduction in calcium signals (baseline, S: 8.3 (2.6), P: 42.3 (14.4); hypoxia, S: 0.1 (0.5), P: 3.0 (3.3). Values are mean (SD) signals/min. Figure 5F). This response was found to be specific for mural cells, since nearby GCaMP6s-labeled astrocytes reacted to the hypoxic conditions with a massive calcium response (Figure 5F).

## Discussion

The present study identifies distinct calcium signatures of different mural cells in the mouse somatosensory cortex. For consistency in the nomenclature of the phenotypically diverse mural cells, we have used the terms smooth muscle cell (SMC), ensheathing pericyte (EP) and capillary pericyte (CP, Grant et al., 2017). These pericyte subtype definitions were already outlined in the earliest descriptions of pericytes almost a century ago (Zimmermann, 1923). Consistent use of these mural cell definitions will improve comparability between studies and lessens possible confusion about findings attributed to these cells.

Recent studies have reported frequent calcium transients in somata and in microdomains along CP processes (Hill et al., 2015, Rungta et al., 2018). We have expanded these observations and found that all mural cells along the arteriovenous axis exhibit basal calcium fluctuations (Figure 2A, Videos 1-4). During spontaneous vasomotor activity, SMCs exhibited calcium oscillations that are inversely related to vessel diameter changes (Figure 2D). These rhythmic pulsations were shown to be based on entrainment by ultra-slow fluctuations (0.1 Hz) in γ-band neuronal activity (Mateo et al., 2017). We observed the same behavior of calcium oscillations in EPs on penetrating arterioles (1^st^ to 4^th^ branch order, Figure 2D). This is in line with the expression of α-smooth muscle actin in mural cells of up to 4^th^ branch order vessels as seen by immunohistochemistry staining and the use of transgenic animals (Hartmann et al., 2015a, Hill et al., 2015). In recent computational and experimental works, these vascular segments have also been shown to be the location of blood flow regulation to supply a particular downstream region (Sweeney et al., 2018, Grubb et al., 2019). It is likely that many findings regarding pericyte control of blood flow were based on experiments focusing on EPs (Peppiatt et al., 2006, Hall et al., 2014). At the capillary level, which we defined as vessels with an average diameter of 4 μm and >4^th^ branch order, CPs exhibit asynchronous calcium signals between somata and processes (Figure 2A, C). In agreement with previous studies, we did not find a correlation between vessel diameter and calcium transients in CPs *in vivo* under healthy conditions (Figure 2D, Hill et al., 2015, Rungta et al., 2018). Nonetheless, we have to note, that two-photon imaging is not suitable for the detection of minute changes in vessel diameter in the expected range of 1-3% for capillaries (Grutzendler and Nedergaard, 2019).

In order to understand the regulation of pericyte calcium signals, we performed pharmacological perturbation experiments in acute brain slices. Surprisingly, the calcium signal frequency in CPs persisted *ex vivo* with no substantial difference from the signal frequency *in vivo* (Figure 3C). This hints at an independent blood flow signal, which is rather influenced by the content of solutes and oxygen, available in excess to the cells in the slice. In contrast, EPs showed a pronounced reduction in their calcium signal frequency *ex vivo* (Figure 3A). The lack of intraluminal pressure in acute brain slices produces a loss in basal tone of arteries and arterioles, causing vascular collapse (Figure 3B, Mishra et al., 2014). This leads to cessation of basal calcium oscillations in SMCs and EPs *ex vivo*. A preconstricting factor is often added to the acute brain slice to generate a vascular tone and maintain calcium oscillations in SMCs (Brown et al., 2002, Filosa et al., 2006). However, we found that preconstriction with a widely used thromboxane A_2_ analog (U46619, 100 nM) caused CPs to form cellular extrusions as well as provoking a large cytosolic calcium increase (Figure 3D, Video 5). We therefore omitted to use any preconstricting agents to focus our investigation on CPs.

Other vasomodulators, such as ATP and Endothelin-1, also caused a massive increase in CP cytosolic calcium levels (Figure 4A-C), as has also been seen in SMCs (Hill-Eubanks et al., 2011). In many cases with Endothelin-1, this calcium increase was accompanied by membrane blebbing and cellular protrusions, comparable to stimulation with the thromboxane A_2_ analog. At the same time the global reactivity of the vasoactive agents caused the vasculature to exert motion. From our observations, we do not exclude the possibility of CPs being able to exert force on their underlying vessel in pathological conditions via cytoskeleton rearrangements within the cell (Fernandez-Klett et al., 2010). A comparable scenario was suggested in Aβ-induced constriction of capillaries in Alzheimer’s disease (Nortley et al., 2019). Moreover, high-power optogenetic stimulation of capillary pericytes produces a similar effect (Hartmann et al., 2020).

Another potent vasomodulator is potassium, which is released in high amounts during the repolarization phase after action potential firing (Paulson and Newman, 1987). Potassium sensing by SMCs and capillary endothelial cells via Kir channels has been previously described as a mechanism to increase local cerebral blood flow (Longden and Nelson, 2015, Longden et al., 2017, Filosa et al., 2006, Haddy et al., 2006). Furthermore, a very recent modeling study showcases the capillary Kir channel as sensor of neuronal signals and highlights its impact on K^+^ - mediated neurovascular communication (Moshkforoush et al., 2020). Based on our experimental data, we suggest that CPs are also able to sense potassium via Kir and K_ATP_ channels (Figure 6): Firstly, we showed that CP calcium signals ceased after excitatory chemogenetic stimulation of neurons *in vivo* (Figure 5A, Video 6). A comparable behavior was found in odorant-stimulated neurons (Rungta et al., 2018) or during spreading depolarization (Khennouf et al., 2018). Secondly, elevation of extracellular potassium *ex vivo* led to a strong reduction in CP calcium signals (Figure 5B) and could be prevented by blocking Kir and K_ATP_ channels (Figure 5C, D), suggesting these channels are involved in silencing the calcium response in CPs. Thirdly, hypoxic insults *in vivo* or chemical ischemia *ex vivo*, leading to a reduced intracellular [ATP]/[ADP] ratio and likely elevated potassium levels due to neuronal depolarization, also resulted in cessation of CP calcium signals activated by Kir and K_ATP_ channels (Figure 5E, F). Activation of Kir and K_ATP_ channels results in CP hyperpolarization, which can be transmitted to the endothelium through endothelial/pericyte gap junctional coupling. Inter-endothelial gap-junctions facilitate further retrograde hyperpolarization to upstream SMCs to induce a relaxation of penetrating arterioles (Longden et al., 2017, Iadecola, 2017). Electrotonic transmission between neighboring pericytes and pericyte/endothelium was demonstrated in the retinal vasculature (Wu et al., 2006) and needs verification in the cortical vasculature. We propose that the mural cell heterogeneity may provide a means of imposing directionality on a propagating hyperpolarization by differential gap-junction coupling with the endothelium to induce hyperemia in a given area. On the other hand, the Kir and K_ATP_-induced hyperpolarization of CPs could also provide a protective mechanism to manage over excitability and lowered metabolic supply during hypoxia/ischemia, as has been proposed for a subset of neurons in the brain (Ballanyi, 2004, Yamada and Inagaki, 2005).

**Figure 6:**
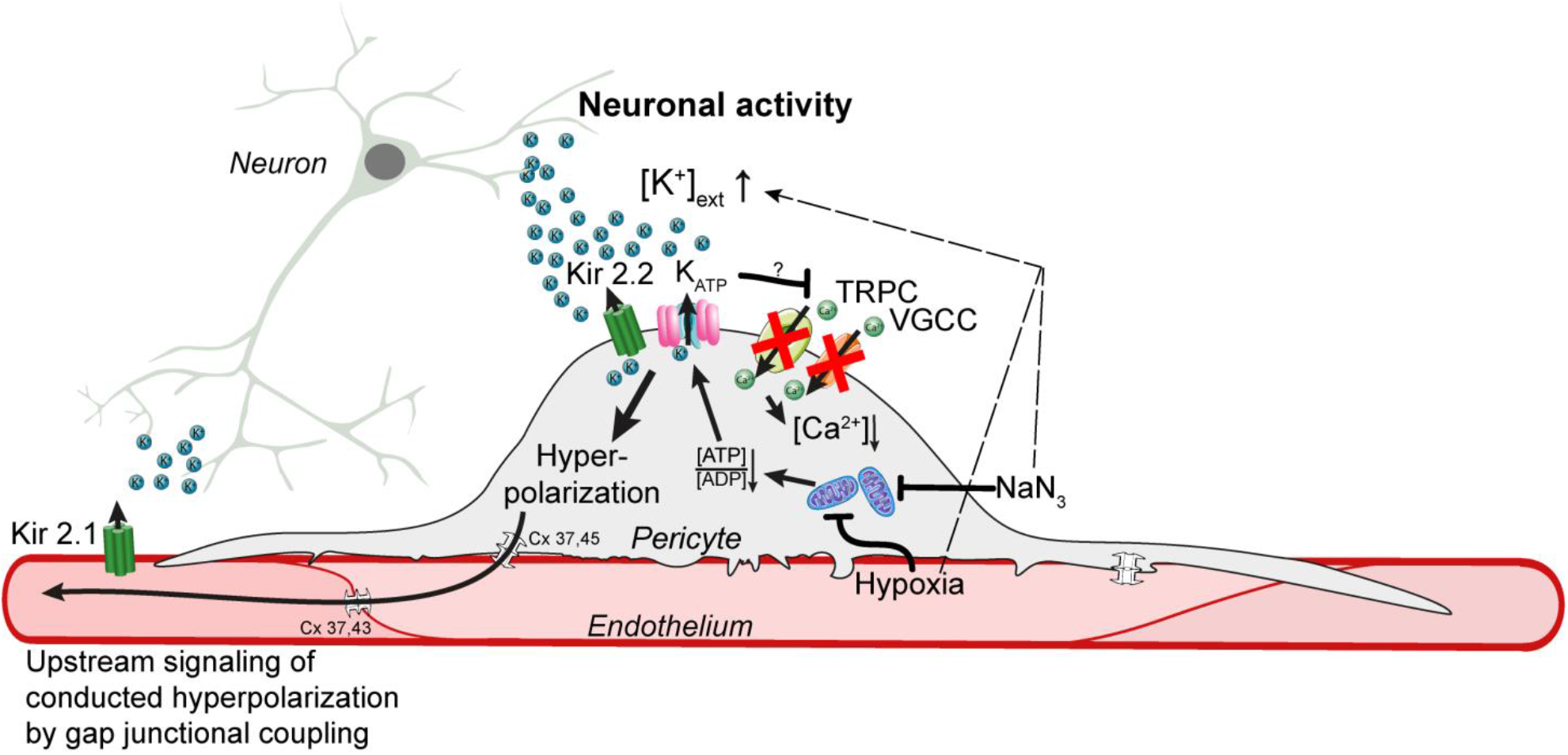
Proposed mechanisms of potassium-induced calcium signal drop in capillary pericytes. Neuronal activity leads to the release of high amounts of potassium into the extracellular space. This rise in potassium activates Kir2.2 and K_ATP_ channels on capillary pericytes to induce a hyperpolarization, which is first transmitted to the gap-junction (Cx 37,45) coupled endothelium and is then further conducted retrogradely between inter-endothelial gap-junctions (Cx 37,43) to upstream SMCs. The pericytic hyperpolarization inactivates TRPC and VGCC channels leading to a drop in calcium signals. Blockage of the respiratory metabolism by NaN3 or hypoxia leads to a presumed reduced [ATP]/[ADP] ratio activating K_ATP_ channels to induce a hyperpolarization and a subsequent decrease in calcium signals.

Calcium is involved in a plethora of cellular processes, ranging from cellular homeostasis to force generation (Berridge et al., 2003). This may explain the spread of calcium signals observed in CPs at basal conditions. The oscillatory nature of the CP calcium dynamic could be a way of reducing the threshold for activation of calcium-related pathways, such as regulation of gene expression (Dolmetsch et al., 1998). Thus, CPs, optimally positioned in the capillary bed at the site of nutrient exchange, may be able to sense slight changes in their environment and adapt their calcium signals accordingly to induce intra and intercellular responses. Further knowledge of the pericyte calcium-signaling toolkit may help to understand the consequences of altered calcium signaling in pericytes in physiological and pathophysiological conditions.

## Materials and Methods

### Animals

All animal experiments were approved by the local Cantonal Veterinary Office in Zürich (license ZH 169/17) and conformed to the guidelines of the Swiss Animal Protection Law, Swiss Veterinary Office, Canton of Zürich (Animal Welfare Act of 16 December 2005 and Animal Protection Ordinance of 23 April 2008). The following mice were interbred: B6.Cg-Tg(*Pdgfrb*-CreERT2)6096Rha/J (*Pdgfrb*-CreERT2), The Jackson Laboratory (Stock #029684, Gerl et al., 2015) with Ai96 (GCaMP6s), The Jackson Laboratory (Stock #028866). Male and female offspring aged 2-9 months were used for experiments. The mice were given free access to water and food and were maintained under an inverted 12/12 hour light/dark cycle.

### Experimental Timeline

To induce GCaMP6s expression, Tamoxifen (Sigma-Aldrich, catalog number T5648), dissolved in corn oil (10 mg/ml), was injected intraperitoneally in mice at a dose of 100 mg/kg on four consecutive days, three weeks before *ex vivo* or *in vivo* imaging. After cranial window implantation, mice were allowed to recover for three weeks prior to *in vivo* imaging, to ensure that all surgery-related inflammation had resolved.

### Anesthesia

For surgery, animals were anesthetized with a mixture of fentanyl (0.05 mg/kg bodyweight; Sintenyl; Sintetica), midazolam (5 mg/kg bodyweight; Dormicum, Roche), and medetomidine (0.5 mg/kg bodyweight; Domitor; Orion Pharma), administered intraperitoneally. Anesthesia was maintained with midazolam (5 mg/kg bodyweight), injected subcutaneously 50 min after induction. To prevent hypoxemia, a face mask provided 300 ml/min of 100% oxygen. During two-photon imaging mice were anesthetized with 1.2% isoflurane (Attane™; Piramal Healthcare, India) and supplied with 300 ml/min of 100% oxygen. Core temperature was kept constant at 37 °C using a homeothermic heating blanket system (Harvard Apparatus) during all surgical and experimental procedures. The animal’s head was fixed in a stereotaxic frame (Kopf Instruments) and the eyes were kept moist with ointment (vitamin A eye cream; Bausch & Lomb).

### Head-post implantation

A bonding agent (Gluma Comfort Bond; Heraeus Kulzer) was applied to the cleaned skull and polymerized with a handheld blue light source (600 mW/cm^2^; Demetron LC). A custom-made aluminum head post was connected with dental cement (Synergy D6 Flow; Coltene/Whaledent AG) to the bonding agent on the skull for later reproducible animal fixation in the microscope setup. The skin lesion was treated with antibiotic ointment (Neomycin, Cicatrex; Janssen-Cilag AG) and closed with acrylic glue (Histoacryl; B. Braun). After surgery the animals were kept warm and were given analgesics (Temgesic™ (buprenorphine) 0.1 mg/kg bodyweight; Sintetica).

### Virus injection and cranial window surgery

A 4 × 4 mm craniotomy was performed above the somatosensory cortex using a dental drill (Bien-Air Dental), and for experiments requiring chemogenetics, adeno-associated virus (AAV) vectors were injected into the primary somatosensory cortex to achieve sensor protein expression: 150 nl of AAV2-hSYN-hM3D (Gq)-mCherry (titer 1.02 × 10^12^ VG/ml; viral vector core facility (VVF), University of Zürich). Large vessels were avoided to prevent bleeding. A square Sapphire glass coverslip (3 × 3 mm, UQG Optics) was placed on the exposed dura mater and fixed to the skull with dental cement, according to published protocols (Holtmaat et al., 2009).

### Two photon imaging

Two-photon imaging was performed using a custom-built two-photon laser scanning microscope (2PLSM) (Mayrhofer et al., 2015) with a tunable pulsed laser (MaiTai HP system, Spectra-Physics and Chameleon Discovery TPC, Coherent Inc.) and equipped with a 20x (W Plan-Apochromat 20x/1.0 NA, Zeiss) or 25x (W Plan-Apochromat 25x/1.05 NA, Olympus) water immersion objective. During *in vivo* measurements, the animals were head-fixed and kept under anesthesia as described above. To visualize the vasculature Texas Red Dextran (2.5% w/v, 70,000 mw, 50μl, Life Technologies, catalog number D-1864) was injected intravenously. GCaMP6s and Texas Red were excited at 940 nm, and emission was detected with GaAsP photomultiplier modules (Hamamatsu Photonics) fitted with 535/50 nm and 607/70 nm band pass filters and separated by a 560 nm dichroic mirror (BrightLine; Semrock). Control of microscope laser scanning was achieved with a customized version of ScanImage (r3.8.1; Janelia Research Campus; Pologruto et al., 2003).

At the beginning of each imaging session, z-stacks of the area of interest were recorded to identify the branch order of individual capillary segments. Once capillaries with a branch order of >4 were identified, a high resolution (512×512 pixels, 0.74 Hz) image was collected for reference and then baseline images (128×128 pixels; 11.84 Hz) were collected over a 90 s period with zoom factors ranging from 12 to 19. Multiple imaging sessions were conducted on different days for each animal.

### Acute brain slice preparation

Prior to slicing, 100 μl of a 2.5%, 70 kDa Texas Red Dextran (Life Technologies catalog number D-1864) was injected intravenously into the tail vein to stain blood plasma. Mice were euthanized by decapitation after deep anesthesia with isoflurane. The brain was extracted from the skull in ice-cold cutting solution consisting of: 65 mM NaCl, 2.5 mM KCl, 0.5 mM CaCl_2_, 7 mM MgCl_2_, 25 mM glucose, 105 mM sucrose, 25 mM NaHCO_3_ and 2.5 mM Na_2_HPO_4_. Slices of 300 μm thickness were cut using a Microm HM 650 V vibratome. Slices were immediately transferred into artificial cerebrospinal fluid (aCSF) consisting of: 126 mM NaCl, 3 mM KCl, 2 mM CaCl_2_, 2 mM MgCl_2_, 25 mM glucose, 26 mM NaHCO_3_ and 1.25 mM NaH_2_PO_4_, at 34 °C. After a recovery phase of 1 hour, slices were used for imaging. All solutions were continuously gassed with 95% O_2_, 5% CO_2_ and were prepared on the day of experiment.

### Pharmacology in brain slices

Slices were imaged at 34 °C in the same aCSF that was used for recovery after cutting. Solutions were infused by gravity into the slice chamber at a rate of 1.5 ml/min with an In-line Heated Perfusion Cube (HPC-G; ALA Scientific Instruments).

Nimodipine (Cat. No. 0600), U46619 (Cat. No. 1932), Endothelin-1 (Cat. No. 1160), TTX citrate (Cat. No. 1069), SKF96365 (Cat. No. 1147) and PNU-37883A HCl (Cat. No. 2095) were obtained from Tocris Bioscience (USA). UDP-Glucose (Cat. No. 94335), ATP (Cat. No. A2383) and all salts were obtained from Sigma-Aldrich.

Nimodipine, SKF96365 and PNU-37883A HCl were dissolved in DMSO and Endothelin-1, UDP-Glucose, ATP and TTX citrate were dissolved in water as stock solutions at the highest possible molarity, stored at −20 °C and were diluted to the desired concentration in aCSF prior to use.

For aCSF with increased potassium concentrations, the sodium ion quantity was adapted to maintain the osmolarity of the solution constant.

### Hypoxia protocol

To induce an acute hypoxic insult to the mice, the oxygen supply was transiently replaced with 2.0 l/min 100% nitrogen for 45 seconds.

### DREADD activation

Neurons expressing hM3D-Gq-DREADD were activated with 30 μg/kg bodyweight clozapine (Tocris, Cat. No. 0444, Jendryka et al., 2019) in saline (0.9% w/v) via a tail vein injection during image acquisition.

### Quantification and statistical analysis

Image analysis was performed using ImageJ and a custom-designed image processing toolbox, Cellular and Hemodynamic Image Processing Suite (CHIPS, Barrett et al., 2018), based on MATLAB (R2017b, MathWorks). For each field of view, all images were spectrally unmixed to reduce potential bleed-through between imaging channels, and aligned using a 2D convolution engine to account for motion and x-y drift in time. Background noise was defined as the bottom first percentile pixel value in each frame and was subtracted from every pixel. Regions of interest (ROIs) were selected by combining 2 distinct methods in CHIPS: hand-selection of cell bodies as well as whole cell; and automated ROI detection with an activity-based algorithm (Ellefsen et al., 2014), both using anatomical images (128 × 128 pixel). A 2D spatial Gaussian filter (σ_xy_ = 2 μm) and a temporal moving average filter (width = 1 s) were applied to all images to reduce noise. A moving threshold for each pixel was defined in the filtered stack as the mean intensity plus 7 times the standard deviation of the same pixel during the preceding 2.46 s. Using this sliding box-car approach, active pixels were identified as those that exceeded the threshold. Active pixels were grouped in space (radius = 2 μm) and time (width = 1 s). Resulting ROIs with an area smaller than 4 μm^2^ were considered to be noise and were excluded. We then combined the previously hand-selected ROIs into a single mask by subtracting the soma ROI from the whole cell territory ROIs, thereby leaving a mask of pericytes without soma. We then multiplied this 2D mask with each frame of the 3D mask obtained from the automated ROI detection in order to obtain a mask of ROIs within the cell territory and outside of the soma. We then extracted traces from two sets of ROIs for each image: the hand-selected soma ROIs and the adjusted 3D activity mask. The minimum distance from each activity ROI to the nearest soma was defined as the shortest distance between ROI edges. The signal vector (dF/F) from each ROI was calculated using the mean intensity between 0.4 s and 4 s as baseline. Short, fast peaks were identified by applying a digital band-pass filter with passband frequencies (f1 = 0.025 Hz and f2 = 0.2 Hz) before running the MATLAB findpeaks function. Noise peaks, due to motion or inflow of high-fluorescent particles were manually removed.

Line scan acquisitions for vessel diameter were also analyzed using our custom-designed image processing toolbox for MATLAB, employing implemented methods described earlier (Kim et al., 2012, Drew et al., 2010, Gao and Drew, 2014).

Statistics for *in vivo* data was performed in RStudio (version 1.0.136) using the lme4 package for linear mixed-effects models (Bates et al., 2015). For fixed effects, we used the experimental condition (with/without drug) or cell type. For random effects, we had intercepts for individual animals and cells. Likelihood ratio tests comparing models with fixed effects against models without fixed effects were used to determine the model with the best fit while accounting for the different degrees of freedom. Visual inspection of residual plots did not reveal any obvious deviations from homoscedasticity or normality. All data were reported and plotted as uncorrected means. P values for different parameter comparisons were obtained using the multcomp package with Tukey post hoc tests.

Graphics and statistical analyses of *ex vivo* experiments were performed using GraphPad Prism (version 7.00; GraphPad Software, La Jolla, CA, USA). Datasets were tested with a Shapiro-Wilk normality test for Gaussian distribution. If one or both of the datasets passed the normality test, a two-tailed paired t-test was carried out; otherwise a Wilcoxon matched-pairs signed rank test was chosen. For multiple comparisons a one-way ANOVA was performed, using the Greenhouse-Geisser correction for sphericity and Tukey post hoc tests for group comparisons. All statistical tests used to evaluate significance are indicated in the Figure legends along with the *P* values. Values for area under the curve (auc) were calculated from 20 s windows of normalized traces for both baseline and drug responses considering the whole cell (soma and processes). For all *in vivo* and *ex vivo* experiments, N gives the number of animals and n is the number of cells. Statistical significances are highlighted based on: \**P* < 0.05, \*\**P* < 0.01, \*\*\**P* < 0.001.

## Data and software availability

The CHIPS toolbox for MATLAB is freely available on GitHub (https://ein-lab.github.io/; Barrett et al., 2018). The source code for calcium signal detection with CHIPS is available as source code file.

## Acknowledgements

We thank the Viral Vector Facility of the University of Zürich for the supply of AAV vectors.

We are grateful to Marc Zünd for assembly and maintenance of two-photon microscopes. Furthermore, we thank Noemi Binini and Zoe Looser for genotyping help. Karen Everett is thanked for critical proofreading of the manuscript.

This project was supported by the Swiss National Science Foundation (Grant 310030_182703 and 31003A_156965).

## Supplementary Figures

**Figure 1 – figure supplement 1:**
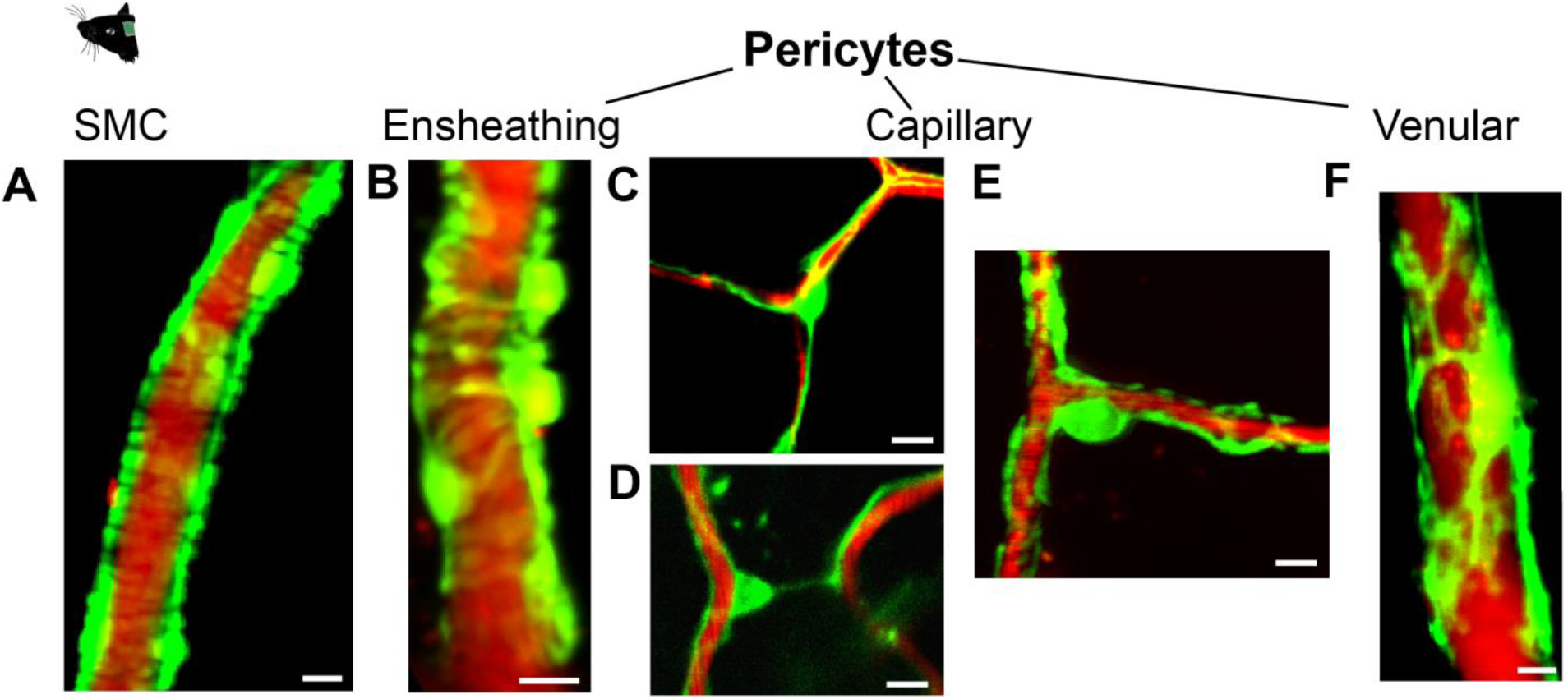
GCaMP6s expression in mural cells of *Pdgfrβ*-CreERT2:R26-GCaMP6s^f/stop/f^ mice. (A)-(F): Distinct morphological appearances of mural cells. Blood plasma is stained with 70 kDa Texas Red Dextran. (A) Ring-shaped smooth muscle cells enwrapping an artery. (B) Ensheathing pericytes enwrapping an arteriole. (C) Thin-strand capillary pericyte on a bifurcation. (D) Thin-strand capillary pericyte interconnecting vessel branches. (E) Mesh pericyte, also categorized as a capillary pericyte, showing net-like process structure. (F) Venular pericyte with stellate morphology on postcapillary venules. Scale bars: 10 μm.

**Figure 1 – figure supplement 2:**
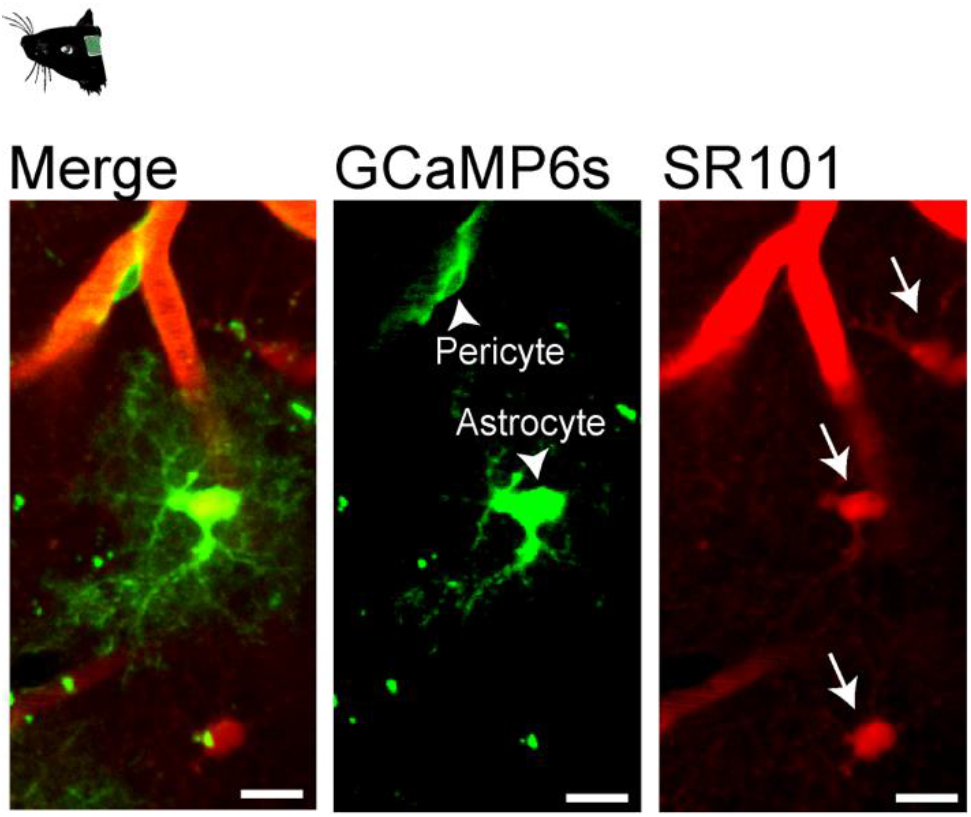
GCaMP6s expression in *Pdgfrβ*-positive astrocytes of *Pdgfrβ*-CreERT2:R26-GCaMP6s^f/stop/f^ mice. Intravenous injection of sulforhodamine 101, SR101 (100 μl of 5 mM iv.) to identify GCaMP6s-expressing astrocytes. Arrowheads point to GCaMP6s-expressing cells in green (GCaMP6s) channel and arrows point to SR101-stained astrocytes in red (SR101) channel. Scale bars: 10 μm.

**Figure 2 – figure supplement 1:**
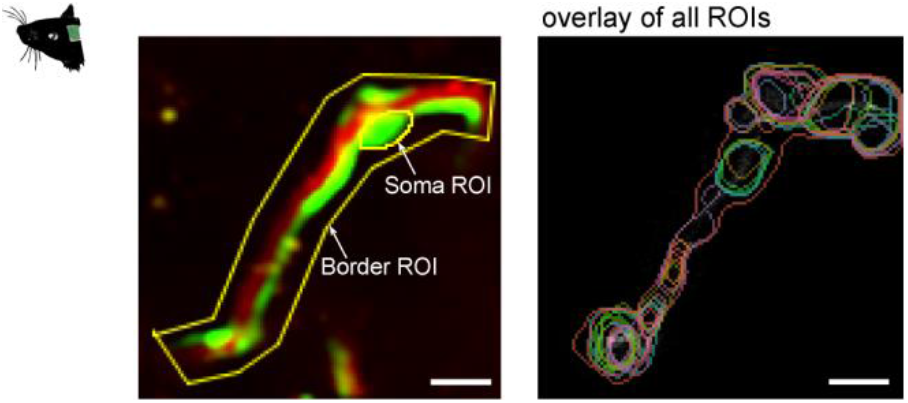
Automatic ROI detection. Example image of ROI selection by hand in ImageJ. The soma was outlined and a border ROI was set around the whole pericyte, so as to reduce detection of signals not associated with the observed cell. To the right is a time overlay of all process ROIs found inside the border ROI. Scale bars: 10 μm.

**Figure 2 – figure supplement 2:**
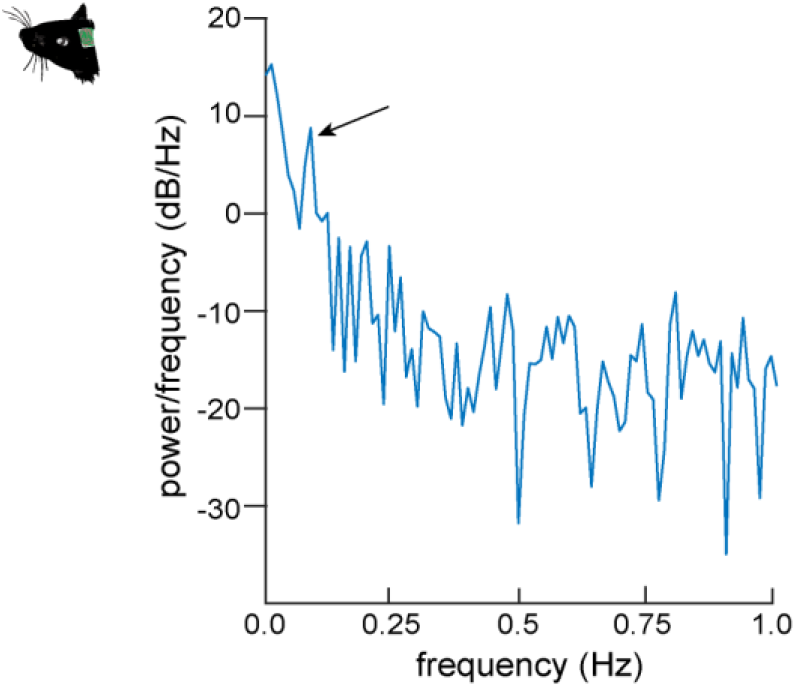
Periodogram of EPs. Periodogram depicting the Fourier transform of the calcium oscillations of EPs in the ultra-low frequency range. The arrow points to a distinct peak at 0.1 Hz.

**Figure 3 – figure supplement 1:**
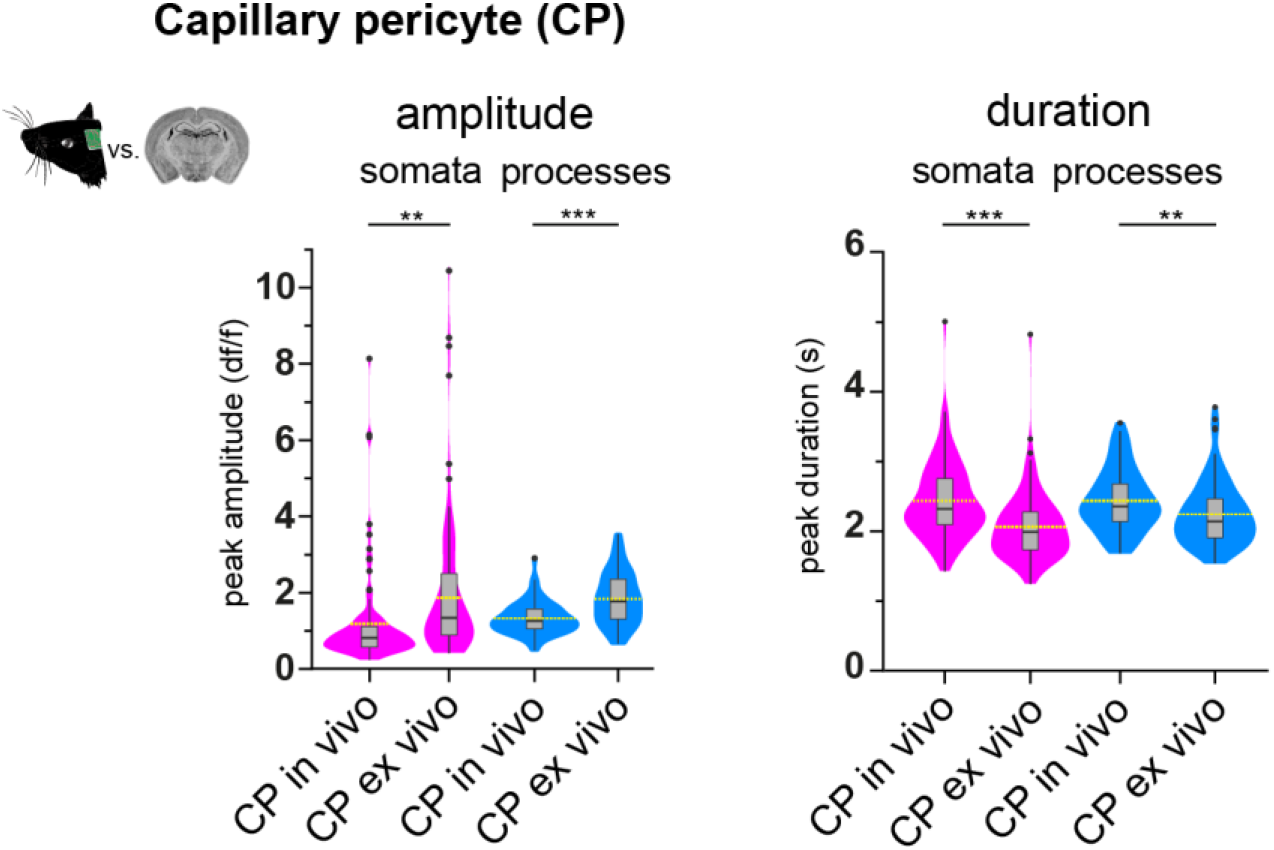
CP calcium signals *in vivo* compared to *ex vivo*. (A) Violin/box plots depicting the quantified calcium signal amplitude and duration comparing capillary pericytes *in vivo* with *ex vivo* for somata and processes. The dashed yellow lines indicate the mean value of the respective parameter. Statistics were calculated using linear mixed-effects models and Tukey post hoc tests. *In vivo*: N=25, n=93, *ex vivo*: N=52, n=106. Amplitude, S: *P* = 0.004, P: *P* = < 0.001; duration, S: *P* < 0.001, P: *P* = 0.006. Source data: Figure 3 – figure supplement 1 – Source Data 1

**Figure 3 – figure supplement 2:**
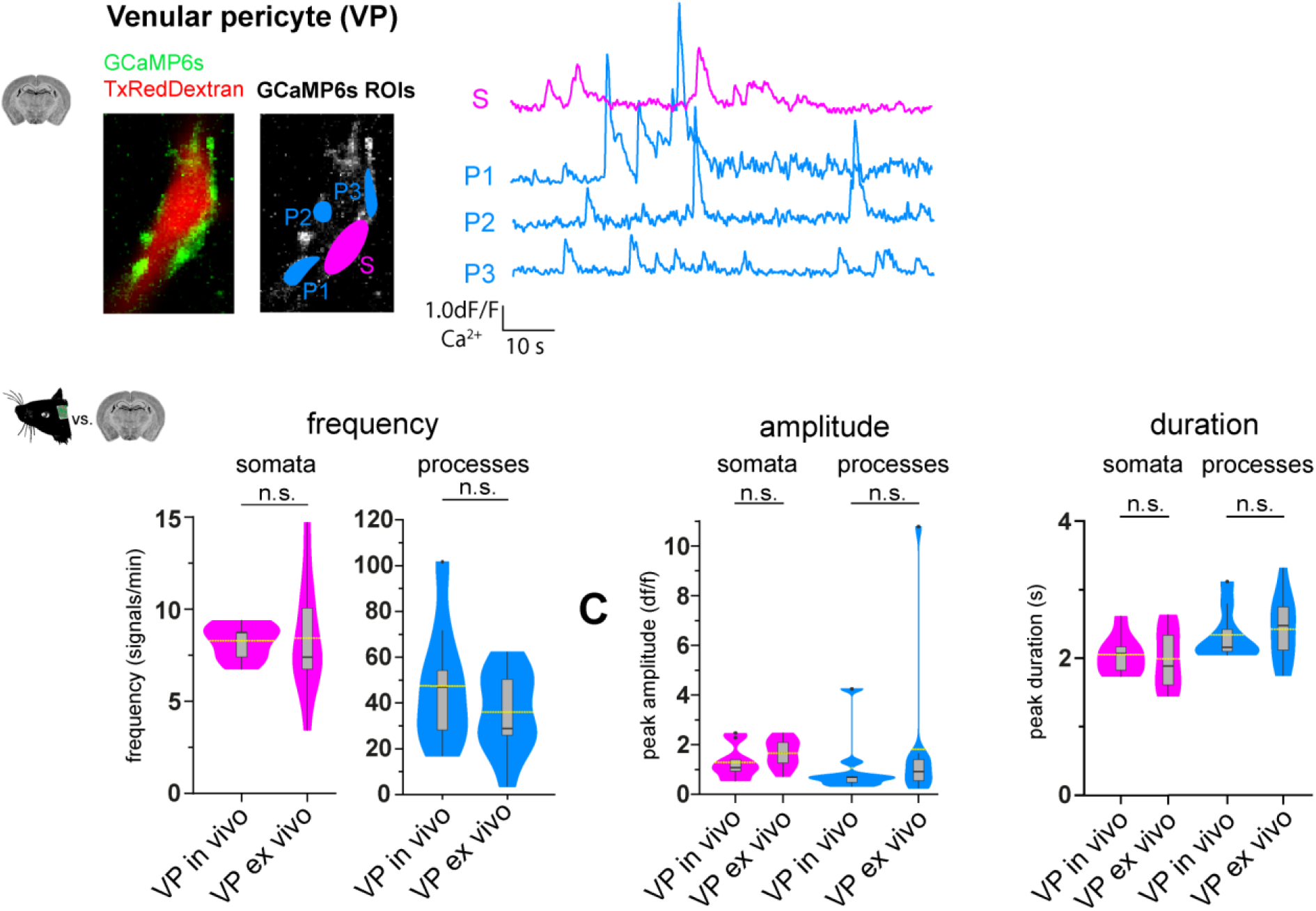
VP calcium signals *in vivo* compared to *ex vivo*. (B) Representative image of a venular pericyte *ex vivo* (left). In the GCaMP6s channel, ROIs for soma (S, in magenta) and processes (P1-3, in blue) are shown. The respective normalized traces of calcium signals are shown to the right. Below, are the violin/box plots depicting the quantified calcium signal frequency, amplitude and duration comparing *in vivo* and *ex vivo* VPs for somata and processes. The dashed yellow lines indicate the mean value of the respective parameter. Statistics were calculated using linear mixed-effects models and Tukey post hoc tests. *In vivo*: N=3, n=9, *ex vivo*: N=6, n=11. Frequency, S: *P* = 0.998, P: *P* = 0.252; amplitude, S: *P* = 0.468, P: *P* = 0.179; duration, S: *P* = 0.723, P: *P* = 0.656. Source data: Figure 3 – figure supplement 2 – Source Data 1

## Video descriptions

### Video 1

Basal SMCs calcium signaling and vasomotion *in vivo*. Recorded at 11.84 Hz and played at 45 fps. Scale bar 5 μm.

### Video 2

Basal ensheathing pericyte calcium signaling and vasomotion *in vivo*. Recorded at 11.84 Hz and played at 45 fps. Scale bar 5 μm.

### Video 3

Basal capillary pericyte calcium signaling *in vivo*. Recorded at 11.84 Hz and played at 45 fps Scale bar 5 μm.

### Video 4

Basal venular pericyte calcium signaling *in vivo*. Recorded at 11.84 Hz and played at 45 fps. Scale bar 5 μm.

### Video 5

*Ex vivo* application of U46619 (100 nM) leads to a massive calcium response in capillary pericytes with cytoplasmic extrusions. For better visibility recorded at 1.5 Hz and played at 20 fps Scale bar 10 μm.

### Video 6

Excitatory chemogenetic activation of neurons leads to a calcium signal drop in pericytes. Recorded at 11.84 Hz and played at 45 fps. Scale bar 10 μm.

## Source data files

Figure 2 – Source Data 1

Figure 3 – Source Data 1

Figure 3 – figure supplement 1 – Source Data 1

Figure 3 – figure supplement 1 – Source Data 2

Figure 4 – Source Data 1

Figure 5 – Source Data 1

Source Code file for CHIPS

